# ABBA, a novel tool for whole-brain mapping, reveals brain-wide differences in immediate early genes induction following learning

**DOI:** 10.1101/2024.09.06.611625

**Authors:** Nicolas Chiaruttini, Carlo Castoldi, Linda Maria Requie, Carmen Camarena-Delgado, Beatrice dal Bianco, Johannes Gräff, Arne Seitz, Bianca A. Silva

## Abstract

Unbiased characterization of whole-brain cytoarchitecture represents an invaluable tool for understanding brain function. For this, precise mapping of histological markers from 2D sections onto 3D brain atlases is pivotal. Here, we present two novel software tools facilitating this process: Aligning Big Brains and Atlases (ABBA), designed to streamline the precise and efficient registration of 2D sections to 3D reference atlases, and BraiAn, an integrated suite for multi-marker automated segmentation, whole-brain statistical analysis, and data visualisation. Combining these tools, we performed a comprehensive comparative study of the whole-brain expression of three of the most widely used immediate early genes (IEGs). Thanks to their neural activity-dependent expression, IEGs have been used for decades as a proxy of neural activity to generate unbiased mapping of activity following behaviour, but their respective induction in response to neuronal activation across the entire brain remains unclear. To address this question, we systematically compared the brain-wide expression cFos, Arc and NPAS4, three abundantly used IEGs, across three different behavioural conditions related to memory. Our results highlight major differences in both their distribution and induction patterns, indicating that they do not represent equivalent markers across brain areas or activity states, but can provide instead complementary information.

## Introduction

The comprehensive understanding of the brain’s functional organisation requires mapping neural circuits at the whole-brain level. Recent advances in histological and imaging techniques have enabled the acquisition of large datasets that scale up to the entire brain and to large experimental groups. This can be achieved through intact-brain imaging using technology such as tissue clearing and light sheet microscopy. However, these approaches are limited by high costs, lengthy antibody incubation time, difficulty in scaling up for large sample sizes, complexity in detecting multiple markers simultaneously and reduced applicability to diverse histological approaches (Renier et al., 2016; Goubran et al., 2019; Tyson et al., 2022; Franceschini et al., 2023). An alternative strategy for whole-brain mapping consists of scaling up to the entire brain classical histological analysis of 2D sections using automated serial microscopy. This approach supports virtually any histological technique, thus overcoming most challenges of intact-brain imaging. The subsequent reconstruction of whole-brain data from serial sections necessitates an automated, time-efficient computational method to register, analyse and visualise 2D-derived data in relation to a common 3D reference atlas. Despite significant progress, this still poses significant challenges. Existing 2D to 3D registration solutions are typically labour intensive, highly demanding in terms of computational resources, require programming proficiency, are limited to the mouse (or rarely rat) reference atlas and to coronal sections, require specific data formats including downsampling, and do not allow the prompt combination of automatic and manual registration options. Lastly, most existing methods do not include a module for segmentation and co-localization of multiple markers and lack an integrated pipeline facilitating data management, atlas navigation and whole-brain data analysis and visualisation (Shamash et al., 2018; Tappan et al., 2019; Yates et al., 2019; Sadeghi et al., 2023). To address these limitations, we developed a scalable pipeline that enables a user-friendly, fine-tuned registration of 2D sectioned brain preparations as well as automated multi-marker quantification and statistical analysis. This novel pipeline includes Aligning Big Brains and Atlases (ABBA), a novel software designed to streamline the registration process, facilitating the precise, rapid and user-friendly mapping of 2D images onto reference atlases. Key distinctive features of ABBA include the interactive graphical user interface (GUI) requiring no coding skills, the integration of multiple atlases (mouse ccfv3, rat, zebrafish, developing mouse, human,..), the compatibility with most imaging file formats without conversion, multi-resolution pyramidal image handling enabling high performance while minimizing resources necessities and seamless integration of deep learning-based automatic registrations (Carey et al., 2023) with manual corrections for the easy handling of damaged tissues.

We then developed Brain Analysis (BraiAn), an integrated suite for quantitative multi-marker image analysis of whole-brain registered datasets. BraiAn is composed of a first module for the standardised and robust segmentation and co-localization of multiple markers within a common coordinate 3D-space as well as dedicated modules for statistical analysis of the obtained datasets. BraiAn offers a versatile infrastructure to facilitate large-scale data analysis and data visualisation, streamlining data management, navigation of atlas parcellation depth, data aggregation (within animals and within groups), normalisation and comprehensive group statistics by partial least square analysis (Krishnan et al., 2011). Importantly, BraiAn analysis modules can be applied to virtually any whole-brain datasets, including the ones derived from 3D cleared brains.

ABBA/BraiAn is thus designed to facilitate brain-wide quantitative analysis of virtually any existing histological method such as immunohistochemistry, in situ hybridization, histochemical stainings (e.g. Golgi staining for dendritic spines, or thioflavin for Aβ deposits), spatial transcriptomics, or viral tracing. Here, we provide an example of the versatile application of ABBA/BraiAn in a comprehensive comparative analysis of three IEGs—cFos, Arc, and NPAS4—across three different behavioural conditions. Brain function underlying behaviour is the result of brain-wide activity changes. In light of this, assessing neural activity modulation across the entire brain is central to decipher network processing underlying behaviour. IEGs reflect recent changes in neuronal activity (Renier et al., 2016; Yap and Greenberg, 2018) and have been used for decades (more than 10’000 papers according to a PubMed search in August 2024 have used IEGs to map neuronal activity) to assess the neural basis of behaviour. Their extensive use in neuroscience research is dictated by several factors: they can be adapted to virtually any behavioural function, grant investigation of deep brain areas, can be easily scaled up to large behavioural groups, allow cell type identification by easy co-labeling, and thus represent a cheap and versatile alternative to more sophisticated methods of neuronal activity mapping. Although classically IEG mapping could only be performed on a handful of brain regions, the recent advent of whole-brain imaging has allowed to scale it to brain-wide networks.

Among the most widely used IEGs one finds cFos, Arc, and NPAS4. cFos (Fos proto-oncogene), the best-known and most extensively used IEG, is a transcription factor that forms the AP-1 complex and plays a crucial role in activity-regulated transcription (Greenberg et al., 1986). Arc (Activity-regulated cytoskeleton-associated protein), unlike most IEGs, is not a transcription factor but a cytosolic protein regulating synaptic function through its association with the cytoskeleton (Lyford et al., 1995). Lastly, NPAS4 (Neuronal PAS Domain Protein 4), a more recently characterised IEG, is an activity modulated transcription factor characterised by its selective expression in neurons and unique induction by neuronal activity (Lin et al., 2008). Despite their well-known molecular differences, their respective induction in response to neuronal activation across the brain remains unclear and they are typically used indiscriminately. A few seminal studies have directly compared IEG induction in targeted brain regions such as the visual cortex or ACC (Hrvatin et al., 2018; Mahringer et al., 2022), and have identified fundamental differences in their stimulus-related induction and cell type specificity. However, a systematic brain-wide comparison between these IEGs is currently lacking. For this, we compared their induction in three different behavioural conditions characterized by increasing salience, namely: mice in their home cage, mice exposed to a novel context and mice exposed to a salient associative learning paradigm. This analysis revealed significant differences in both their baseline brain-wide expression as well as their induction following behaviour. In addition, we found that although the fraction of neurons co-expressing multiple IEGs was surprisingly low, co-expression probability increased with behavioural salience in a specific set of hippocampal, cortical and amygdalar regions. This suggests that different IEGs do not represent equivalent markers across brain areas or activity states, but multiplexed IEGs analysis instead can provide complementary information.

## Results

### Aligning Big Brains and Atlases (ABBA), a scalable tool for brain-wide registration of 2D sections

Aligning 2D sections to 3D volumetric atlases is a crucial first step for whole-brain histological quantitative analysis. Although powerful, existing 2D to 3D registration tools still present limitations that hinder their widespread and effortless application. To overcome these limitations, we developed ABBA, an ImageJ/Fiji (Schindelin et al., 2012) -based software, that is the result of the authors’ collective expertise, combining insights from experimental neuroscientists and microscopy core facility staff. We identified technical and practical criteria aimed at offering flexibility, efficiency, ease of use, and reliable results, with user expertise remaining paramount. To start a registration project on ABBA, a reference 3D atlas and a set of experimental slices are selected (Fig. 1A, S1A). For reference atlases, ABBA can load the Allen mouse brain atlas CCFv3 (Wang et al., 2020) or the Waxholm Rat atlas v4 (Papp et al., 2014; Kleven et al., 2023) or any other BrainGlobe compatible atlas (Claudi et al., 2020). Loading and displaying experimental image files represents a limiting factor in existing registration tools as they lack an optimised support for large 2D images. Most existing tools require specific file formats like CZI or MRXS or necessitate downscaled versions of the original data (Shamash et al., 2018; Tappan et al., 2019; Yates et al., 2019; Sadeghi et al., 2023). To address these limitations, ABBA exploits multiresolution file reading by integrating Bio-Formats (Linkert et al., 2010) and BigDataViewer (Pietzsch et al., 2015), which ensure the high processing speed required for an interactive display of a large dataset of high-resolution images (e.g. a typical whole-brain dataset from one mouse is composed of tens of images each typically encompassing gigapixels across multiple channels). Accordingly, we designed ABBA to process a wide range of pyramidal image files typically outputted by slide imagers or shared data-bases (e.g. .svi, .czi, .bif, .dicom, .ome.tif …) without the need for prior conversion or downsampling. Pyramidal image files contain high-resolution as well as pre-computed downsampled image versions which allows ABBA to work at the resolution level needed at any point, thus significantly accelerating the visualisation and processing steps (Fig. S1B).

**Figure 1:**
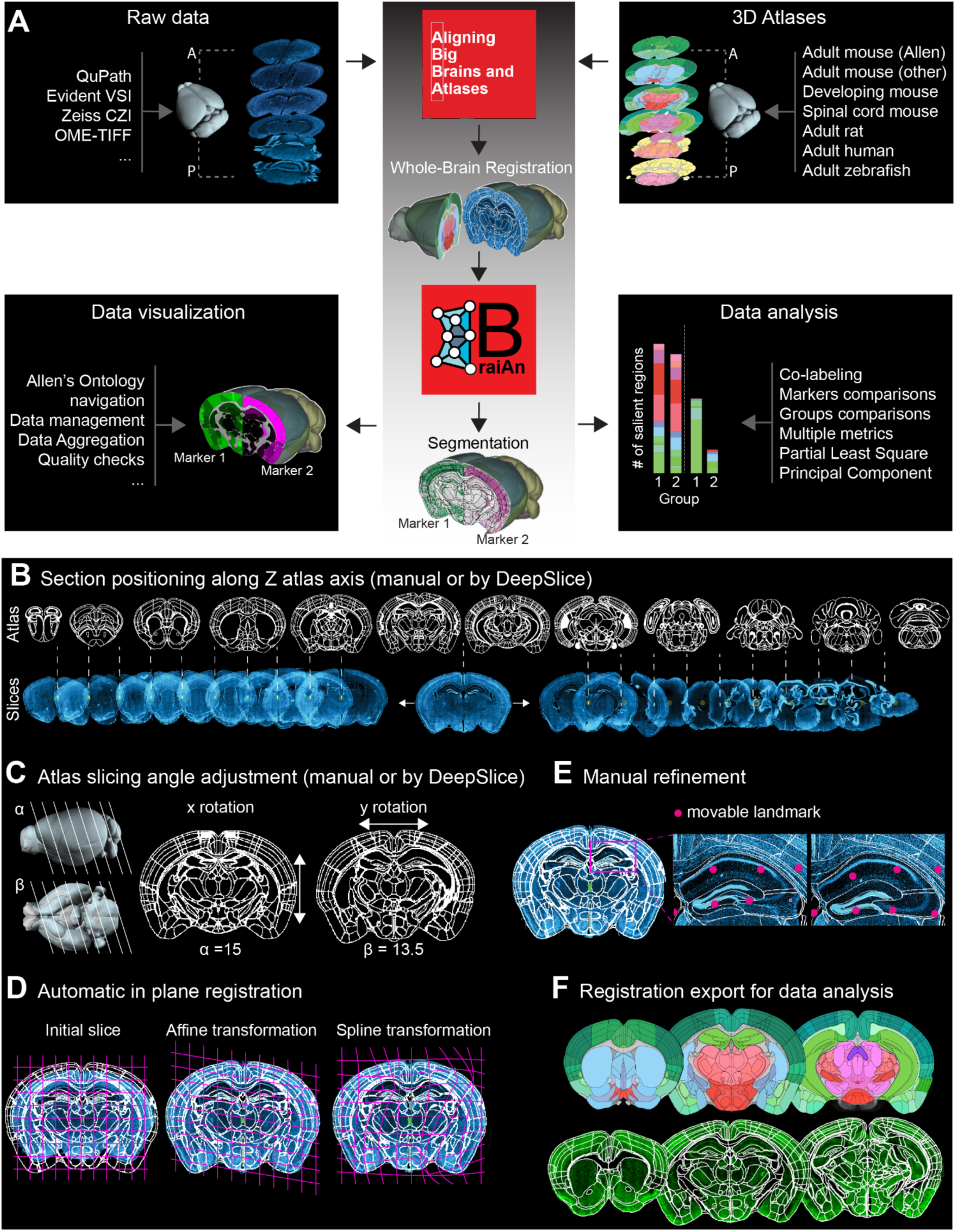
Workflow overview. **A.** ABBA has two inputs: i) a dataset of experimental brain section images, typically packaged as a QuPath project, and ii) a 3D atlas chosen within the mouse CCFv3, rat Waxholm v4, or any other BrainGlobe compatible atlas (Claudi et al., 2020). The output of ABBA’s registration process is the transformation function that maps pixel coordinates from the atlas to the sections, and vice versa. Whole brain registration deformation functions can be directly imported within QuPath, a popular and extensible image analysis software, and, through BraiAnDetect extension, automatic cell segmentation and marker co-expression are quantified for each brain section. Obtained whole-brain registered histological data are analysed and visualised with BraiAn, a versatile Python library (Fig. S1). **B-F.** Main steps of whole brain registration in ABBA: **B.** Experimental sections are positioned along the atlas axis in ABBA. The Z axis is defined according to the atlas cutting orientation (coronal, sagittal, horizontal). Z positioning can be done manually or automatically by direct integration of DeepSlice. **C.** Fine tuning of the atlas cutting angle in 2 orientations (manually or automatically through DeepSlice integration). **D.** In-plane registration: linear (affine) and non-linear (spline) automatic transformations are iteratively performed until optimal alignment is achieved. Further registration refinement is obtained by manual optimization through BigWarp (**E**). **F.** ABBA’s output (a function mapping atlas coordinates to experimental section pixel coordinates) is used to export the transformed atlas data to image analysis software such as QuPath.

After selecting and loading atlas and experimental files, the first step for registration consists of positioning each experimental image along the atlas axis. For this, ABBA first virtually sections the chosen 3D atlas at any cutting angle and displays the obtained atlas sections in the GUI (Fig. 1B,C, S1C, D) with customizable atlas channel visualisation (Nissl, autofluorescence, region outlines) and customizable spacing between sections (Fig. S1C, D). Atlas data consist of the combination of raster 3D multi-channel pixel data (with one channel specifying atlas region index) and a tree ontology specifying the hierarchy of the different regions. The interactive 2D display of 3D atlases derived from this data combination was built exploiting BigDataViewer (Pietzsch et al., 2015) interface with imgLib2 (Pietzsch et al., 2012), which allows the transparent processing of multidimensional image files with enhanced flexibility and efficiency. All experimental sections are then loaded and displayed below (with customizable multi-channel display, Fig. S1D). Section positioning to the matching atlas slice can be manually or automatically performed. For manual positioning, the graphical user interface facilitates convenient movement of sections along the atlas axis and offers the possibility to carefully match a few sections and interpolate the position of intermediate slices assuming evenly spaced cutting, thus significantly accelerating the process. Automating "Z" positioning and atlas cutting angle selection remains challenging. However, recent tools have shown some success. DeepSlice (Carey et al., 2023), for instance, uses machine learning to position slices in the coronal orientation of the mouse brain CCFv3 and rat atlas. Consequently, ABBA directly interacts with DeepSlice for machine-learning based atlas sections and cutting angle matching within ABBA. Importantly, the obtained outputs can be easily manually refined in ABBA, a step not promptly available within DeepSlice. The integration of DeepSlice within ABBA offers an ideal option for rapid, user-friendly positioning and ensures accessibility to users with limited knowledge of brain anatomy. Once each section is correctly positioned along the atlas Z axis, ABBA can perform automatic ‘in-plane’ registrations by adapting the experimental sections to precisely match the atlas coordinates. This is achieved via a concatenation of linear and non-linear transformations. For these automated registration steps, ABBA uses Elastix (Klein et al., 2010) with pre-defined registration parameters and capitalises of its access to the sections’ calibration as well as the atlas physical voxel size in order to have an almost parameter-free registration. The metric used to measure the ‘distance’ between fixed (atlas) and moving (experimental sections) images is the Mattes mutual information metric (Mattes et al., 2003), which enables registering different imaging modalities. Importantly, experimental to atlas section automatic registrations in ABBA are performed using multiple channels. For instance, a nuclear channel (i.e., DAPI) and an autofluorescence channel can be paired with the Nissl and ARA channels of the Allen Brain Atlas respectively. Both pairs of images will be weighted equally in the metric measurement. Importantly, automatic registrations of all slices can be performed in parallel (one registration per CPU core) and do not block enqueuing of other ABBA functions, thus significantly enhancing processing speed (i.e., 3 iterative registrations of 80 brain sections are computed in around 10 minutes with a regular 8-core CPU, Fig. S1E). ABBA also provides an option for manual in-plane registrations by integrating BigWarp (Bogovic et al., 2016). An effective approach consists in a first round of iterative automated linear (affine) and non-linear (spline) registrations, followed by a manual refinement of the output with BigWarp (Fig. 1E). In practice, we have found that combining automated and manual correction of registration yields the most efficient and accurate outcomes. ABBA’s iterative registration strategy provides users with the flexibility to set the balance between registration speed and accuracy for each project, spanning from a rough registration that can be achieved in a few minutes per animal, up to any desired accuracy with increasing time dedicated to alignment refinements.

The output of the registration is, for each slice, an invertible function that maps 3D atlas coordinates to the experimental image pixel coordinates that can be directly used for image analysis in Java or in Python by using PyImageJ (Rueden et al., 2022). However, to facilitate the following image analysis steps, ABBA contains built-in commands for exporting all atlas registered regions as annotations into the popular, flexible and efficient QuPath software (Bankhead et al. 2017) which was designed to work with large datasets of tiled images (Fig. 1F). As a result, quantitative image analysis of ABBA-registered datasets can rely on the wide array of built-in scriptable image analysis tools offered by QuPath including automated object segmentation and interactive machine learning for pixel and object classification. ABBA’s compatibility with QuPath (Chiaruttini et al., 2022), is a crucial asset as it ensures flexibility and consistency in the post-registration analysis steps. In conclusion, ABBA provides a scalable solution for brain-wide quantification of 2D sections, offering flexibility, ease of use, and reliable results while minimising resources requirements. Integrated seamlessly with existing bio-imaging software, ABBA streamlines the alignment process, enabling efficient and accurate whole-brain histological analysis of large groups of animals.

### Brain Analysis (BraiAn), a versatile toolkit for whole-brain quantitative analysis of large datasets

Following the whole-brain 2D to 3D atlas registration provided by ABBA, we developed BraiAn, an open-source suite of tools designed to simplify signal quantification, analysis and visualisation of large datasets typically obtained in whole-brain imaging experiments (Fig. 1A, S2). The first module, named BraiAnDetect, consists of a QuPath extension for multichannel cell segmentation across large and variable datasets ensuring consistency in segmentation settings. This module leverages QuPath’s built-in algorithms to provide a multi-channel, whole-brain optimised cell detection pipeline that runs in a fully automated manner in less than 40 minutes per brain on a regular desktop computer. BraiAnDetect features additional options for refining signal quantification, including machine learning-based object classification, region specific cell segmentation, multiple marker co-expression analysis and an interface for selective exclusion of damaged tissue portions. The second module, named BraiAnalyse, is a modular Python library for the easy navigation, visualisation, and analysis of whole-brain quantification outputs. BraiAnalyse’s workflow chains iterative data aggregation operations from single sections to single animals and single groups (Fig. 1A, S2A) while consistently attributing the resulting data to the atlas ontology and minimising the possibility of mistakes by enforcing type & data checks. The major BraiAnalyse functions include:

1. Guided whole-brain parcellation at different granularities, enabling the selection of non-overlapping regions covering the whole brain. The granularity level can be set based on several parcellation methods including depth in the Allen brain atlas ontology, structural level, Allen’s major divisions and summary structures (Wang et al., 2020), custom…), which is facilitated by an interactive visualisation of the atlas ontology (Fig. S2B).
2. Computation of several normalised per-region metrics, including positive cell density, relative fraction, multiple markers co-localization indices, relative marker changes.
3. The generation of an interactive *XMasTree* plot displaying results of the chosen metrics for all brain regions organised by anatomical major divisions, along with per-subject and per-group scatter plots enabling to easily navigate large whole brain datasets (Fig. S2C, Fig. S3-5).
4. Diverse options for intuitive data visualisation on whole-brain 2D sections representation (Fig. S2B, C), leveraging BrainGlobe (Claudi et al., 2020). Moreover, this library offers options for data quality check, such as systematic evaluations of the inter-sections and inter-animals variability.
5. Lastly, it includes a module for statistical comparisons across experimental groups, using an optimised implementation of task mean-centered partial least squares correlational analysis (Krishnan et al., 2011) (PLS, for implementation details see *Methods*). PLS results are integrated into the *XMasTree* plot and can be visualised in ad hoc plots (Fig S2D-G).

Several features make BraiAnalyse a particularly handy tool for whole-brain analysis: i) it integrates the Allen Brain Atlas hierarchical organisation allowing data visualisation and data analysis at any preferred level of parcellation; ii) it is compatible with any whole-brain data source (not only ABBA); iii) it is open source, fully documented and its implementation requires only a basic programming expertise.

### Brain-wide comparison of cFos and Arc expression

We then exploited the unique capacity of ABBA/BraiAn to perform fast and precise whole brain co-expression analysis of multiple markers, to address the unresolved question of whether different IEGs are equally represented across the mouse brain. For this, we first characterised the brain-wide (co)-expression of the two most widely used IEGs, cFos and Arc. We exposed mice to a mildly salient experience, specifically, a 5-minute exploration of a new context (CTX), and subsequently performed immunohistochemistry (IHC) for the two markers. Using ABBA, we automatically registered images (acquired with a slide scanner fluorescence microscope) to the Allen brain atlas CCFv3 and subsequently manually refined the alignment so that the registration precision would enable the reliable identification of small nuclei and cortical layers. We then used BraiAn for the machine learning-based segmentation of positive cells and the comprehensive analysis of cFos and Arc densities and co-expression across the brain (Fig. 2A). Whole-brain analysis of both markers (Fig. 2A, Fig. S3, 4) revealed striking differences in their distribution. While the density of Arc^+^ neurons is considerably higher than that of cFos^+^ neurons in cortical regions (Fig. 2B, C), subcortical regions including the pallidum, thalamus, hypothalamus, midbrain, pons and medulla show high cFos density but undetectable Arc levels. Within this set of subcortical divisions, we found detectable Arc signal in only two specific regions, in the ventromedial hypothalamus (VMH) and in the reticular nucleus of the thalamus (RT), suggesting that these two nuclei may have distinct transcriptional features as compared to their neighbouring structures. Lastly, only a small subset of divisions, namely, olfactory areas, the hippocampus, cortical subplate and the striatum show comparable density of both markers (Fig. 2B, C).

**Figure 2:**
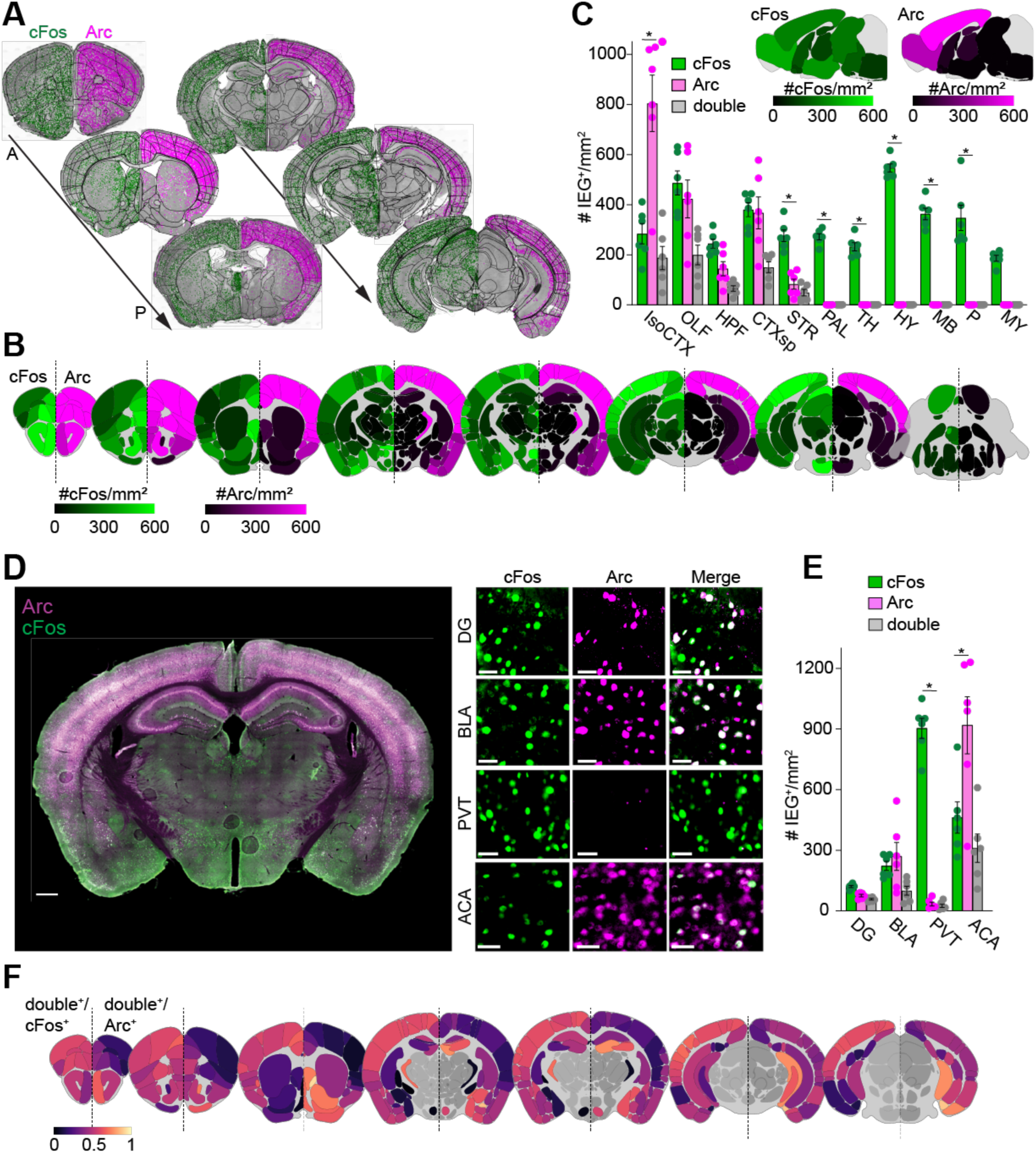
cFos and Arc whole-brain distribution. **A.** Representative examples of atlas registration and positive cell segmentation. Arc detections are shown on the right hemisphere in magenta; cFos ones on the left in green; double positive detections are white. The black reticulate represents the boundaries of Allen CCFv3’s brain regions, after being mapped onto each brain slice with ABBA. **B.** Heatmaps of the whole-brain cFos (left) and Arc (right) densities (mean, N=6 animals). Parcellation level: *summary structures* as defined by CCFv3 (Wang et al., 2020). **C.** cFos, Arc and double positive densities in CCFv3’s *major divisions* (Wang et al., 2020), except for cerebellum (mean ± SEM, N=6 animals): cFos vs Arc two-way ANOVA, F_10,110_ = 17.96, P<0.0001, multiple comparison, Sidak, * P<0.05 **D.** Representative images of cFos and Arc immunohistochemistry. Left: whole-section representation, scale bar 500 µm, right: representative regions: dentate gyrus (DG); basolateral amygdalar (BLA); paraventricular nucleus of the thalamus (PVT) and anterior cingulate area (ACA); scale bar 40 µm. **E.** cFos, Arc and double positive densities in DG, BLA, PVT and ACA (mean ± SEM, N=6): Arc vs cFos two-way ANOVA, F_3,40_ = 36.41, P<0.001, multiple comparison, Sidak, * P<0.05 **F.** Heatmaps of Arc (left) and cFos (right) overlap proportion (mean) of summary structures in Isocortex, OLF, HPF, CTXsp and STR.

We then further utilised the BraiAn module for multiple markers overlap analysis within the brain areas in which detectable expression was present for both, and thereby built a brain-wide IEGs coexpression map. Surprisingly, we found that even in regions in which both markers are highly expressed, their co-expression at the cellular level is relatively low (Fig. 2D-F). Across the brain, the low degree of overlap reflects different conditions. In some areas, such as thalamic and hypothalamic regions, low overlap reflects the presence of only one marker (cFos), while in cortical areas, it reflects the high abundance of Arc over cFos. However, in other areas commonly investigated via IEGs, such as the hippocampal area DG and the amygdala, the two markers have similar densities, but double positive neurons never exceed 50% (Fig. 2D-F) indicating that cFos and Arc are likely marking different neuronal populations across the brain.

### Brain-wide comparison of cFos and NPAS4 expression

We then analysed a third immediate early gene, NPAS4. This gene exhibits several distinctive features from other IEGs: it is selectively expressed in neurons, uniquely activated by neuronal activity rather than by extracellular signals like growth factors and neurotrophins and directly regulates the expression of a significant number of activity-dependent genes (Lin et al., 2008; Ramamoorthi et al., 2011). However, its representation at the whole brain level and the direct comparison with other more classically used IEGs remain unclear. To address this question, we directly compared the whole-brain distribution of NPAS4 to that of cFos in the same novel context-exposed animals used for cFos-Arc comparative analysis (Fig. 2), taking advantage of the possibility to work on parallel section series offered by our 2D to 3D approach. The experimental and analytical workflow used was identical to the one used for the cFos and Arc comparison. Interestingly, across the entire brain, the density of NPAS4 was around one order of magnitude smaller than that of cFos (Fig. 3A-C). However, similarly to what we observed for Arc, its distribution across the brain differed substantially from the relatively uniform brain-wide cFos expression, with enrichment in cortical and hippocampal structures and undetectable expression in most subcortical regions, including the striatal pallidum, thalamus, hypothalamus, midbrain, pons, and medulla (Fig. 3A-C). We then analysed the co-expression between NPAS4 and cFos within the brain areas in which detectable expression was present for both. Interestingly, we found that although NPAS4 expressing cells are drastically lower in number, they were almost exclusively cFos^+^, indicating that NPAS4 represents a small subset of the larger cFos^+^ population. In contrast, only very few cells exclusively expressing NPAS4 were detected (Fig. 3D-F).

**Figure 3:**
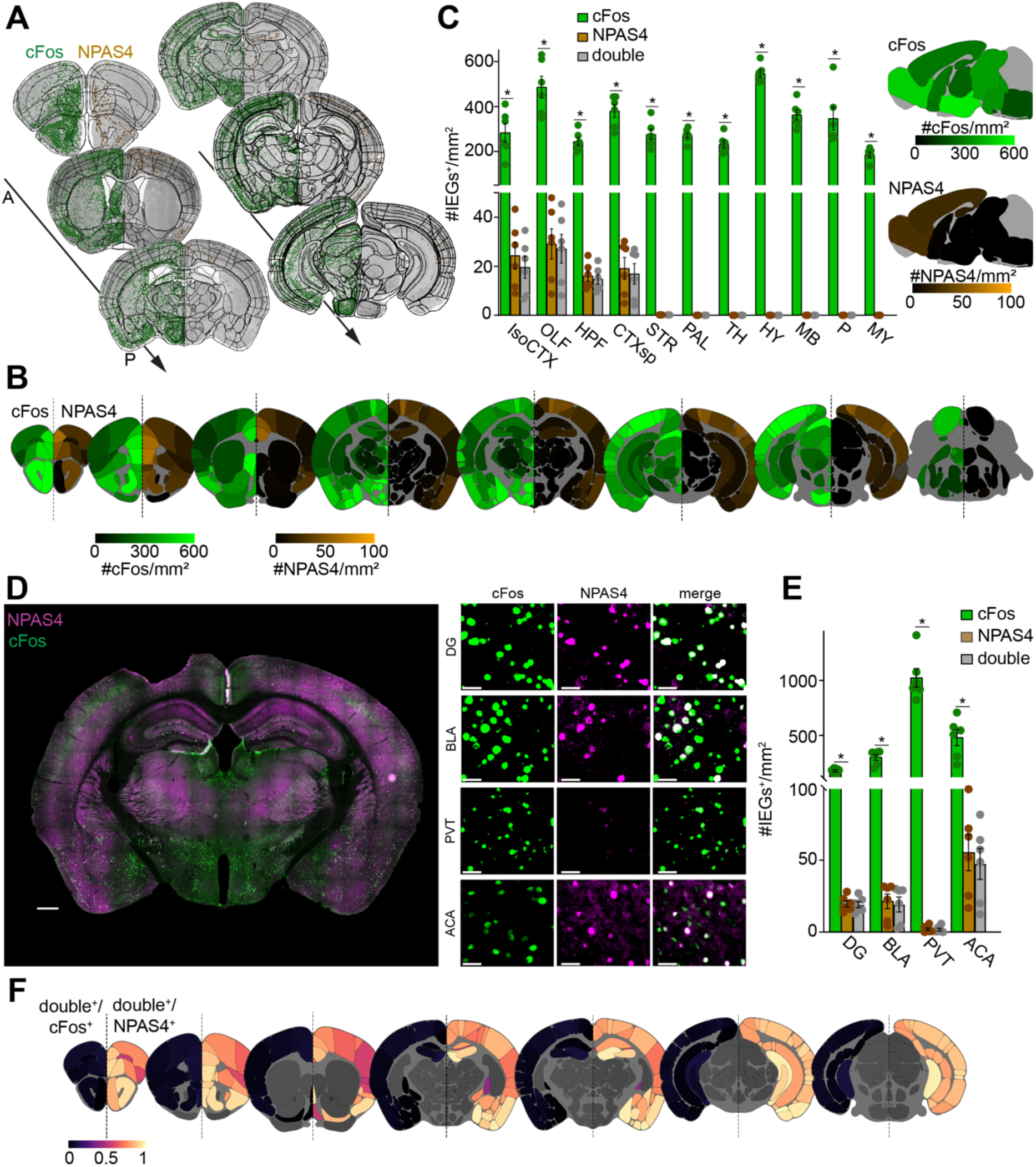
cFos and NPAS4 whole-brain distribution. **A.** Representative examples of atlas annotation and positive cell segmentation. NPAS4 detections are shown on the right hemisphere in brown colour; cFos ones on the left in green; The black reticulate represents the boundaries of Allen CCFv3’s brain regions after being mapped onto each brain slice with ABBA. **B** Heatmaps of the whole-brain cFos (left) and NPAS4 (right) densities (mean, N=6 animals). Parcellation level: *summary structures* as defined by CCFv3 (Y. Wang et al., 2020). **C.** cFos, NPAS4 and double positive densities in CCFv3’s *major divisions* (Y. Wang et al., 2020), except for cerebellum. cFos vs NPAS4 two-way ANOVA, F_10,110_ =13.29, P<0.001, multiple comparison, Sidak, * P<0.05. (Mean ± SEM, N=6 animals). **D.** Representative images of cFos and NPAS4 immunohistochemistry. Right: whole-secction representation, scale bar 500 µm. Left: representative sample regions: DG; BLA; PVT and ACA; scale bar 40 µm. **E.** cFos, NPAS4 and double positive densities in DG, BLA, PVT and ACA. cFos vs NPAS4 two-way ANOVA, F_3,40_ = 43.86, P<0.001. Multiple comparison, Sidak, * P<0.05. Mean ± SEM, N=6 animals. **F.** Heatmaps of NPAS4 (left) and cFos (right) overlap proportion (mean) of summary structures in the isocortex, OLF, HPF and CTXsp.

### Differential brain-wide induction of cFos, Arc and NPAS4 following behaviour

We then investigated whether cFos, Arc and NPAS4 are differentially induced following behaviour. For this, we compared expression patterns between three different experimental conditions. We added to the group exposed to a novel context (CTX), a group subjected to a contextual fear conditioning (CFC) session, which was identical to the novel context experience but with exposure to electrical foot shocks (3 shocks, 0.8 mA, 2s). Undisturbed animals from their home cages were used as baseline controls (Fig. 4A). High resolution, high throughput quantification of the brain wide densities for cFos, Arc and NPAS4 positive cells were obtained thanks to ABBA/BraiAn, which allowed scaling up brain-wide multi-marker analysis to behaviourally relevant group sizes (N=6 animals per group, 3 behavioural groups, 2 series per brain. Fig. S3-5). After having segmented positive cells for the three markers across the brain *summary structures* (as defined by CCFv3), we performed brain-wide global comparisons of their densities between behavioural groups using BraiAn built-in integrated mean-centered task PLS analysis. Interestingly, we found that cFos and NPAS4 brain-wide distributions can efficiently differentiate between all three behavioural groups, with home cage (HC) and FC groups being the most distant and CTX displaying an intermediate brain wide expression pattern (Fig. 4B). In comparison, Arc expression outperforms cFos and NPAS4 in separating the HC group from animals exposed to a novel context but fails in distinguishing CTX from FC groups (Fig. 4B, Fig. S3-5).

**Figure 4:**
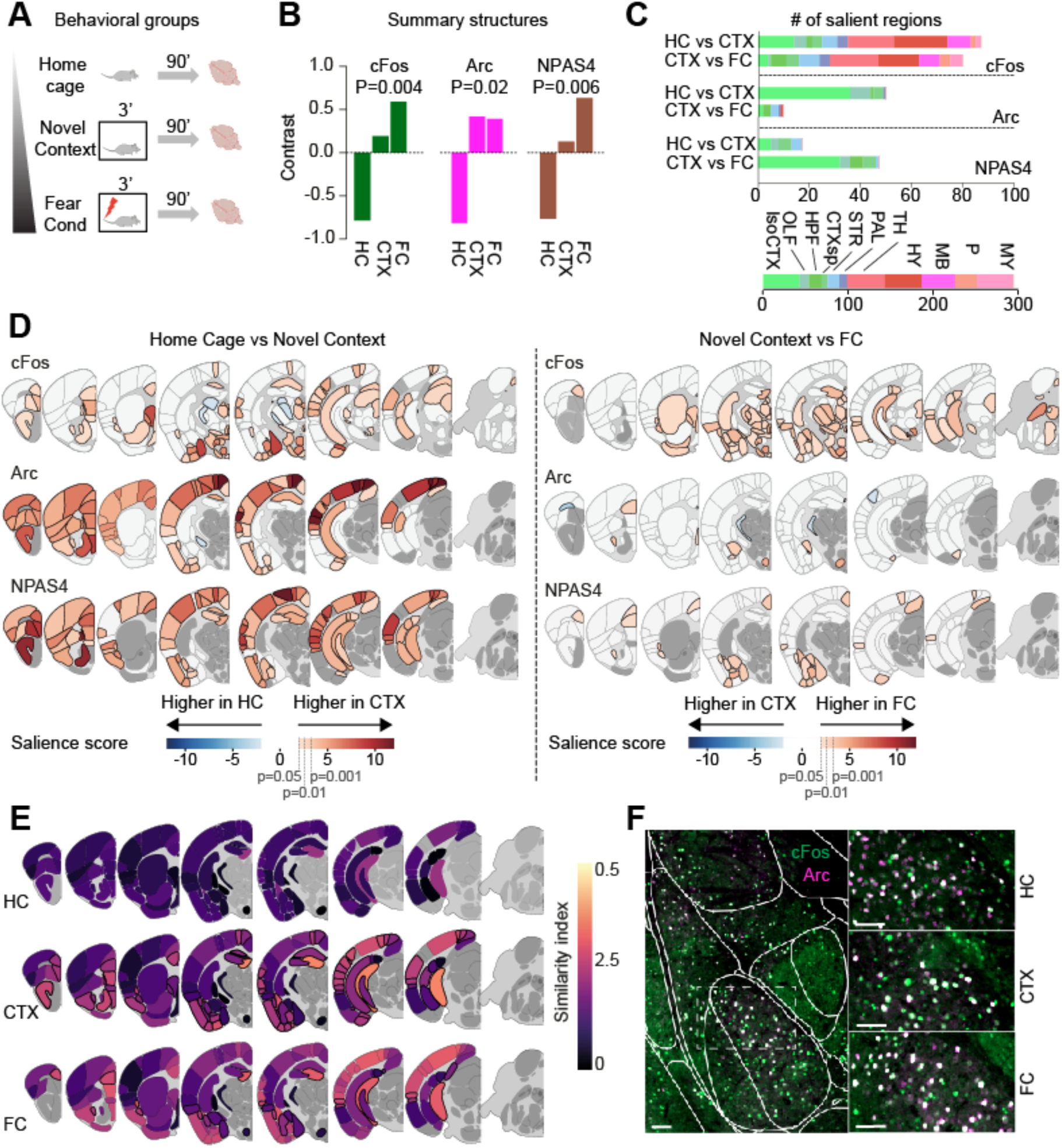
cFos, Arc and NPAS4 induction in response to behavioural stimuli. **A.** Experimental design for IEG analysis. Three behavioural groups were used: animals left undisturbed in their home cages (HC), animals exposed to a novel context for five minutes (CTX), animals exposed to a contextual fear conditioning (FC) session that was identical to the novel context experience but with exposure to electrical foot shocks (3 shocks, 0.8 mA, 2s). **B.** Contrasts of the first latent variable of the mean-centered task PLS analysis between all three groups’ cFos, Arc and NPAS4 density. PLS’s latent variables are generalised with a permutation test (n=10000) and normalised by standard deviation calculated with bootstrap (n=10000). The final score is related to the first left singular vector that maximally differentiates between conditions. Animals, N=6 per group (except for NPAS4, home cage: N=5); Analysed regions (summary structures): cFos: n=206; Arc: n=83, NPAS4: n=60 **C.** (Top) Number of salient summary structures for each of the major divisions in six mean-centered task PLS analyses comparisons: HC vs CTX and CTX vs FC, for cFos, Arc and NPAS4. Salience scores identify regions that differentiate between two conditions by marker density; a region is salient when its score is >1.96 (p<0.05). **D.** Heatmaps of cFos, Arc and NPAS4 density salience scores in the summary structures when comparing HC vs CTX (left), and CTX vs FC (right). If the PLS salience score of a brain region is not significant (≤1.96), it is white-coloured. For cFos CB was excluded. For Arc CB, TH, HY, MB, P and MY were excluded because no detectable expression was found. For NPAS4 CB, TH, HY, MB, P, MY, STRd, STRv and LSX were excluded because no detectable expression was found. **E.** Heatmaps of cFos and Arc *similarity index* (mean) of summary structures in Isocortex, OLF, HPF, CTXsp and STR, in all three conditions. Regions outlined by a black border in CTX indicate the structures whose similarity index changes significantly compared to Home Cage (score>1.96). Regions outlined by a black border in FC heatmaps, indicate significant differences compared to the CTX group (score>1.96). **F.** Representative images of cFos and Arc immunohistochemistry. Left, representative amygdalar region. The white reticulate represents the boundaries of Allen CCFv3’s *leaf regions*, after being mapped onto each brain slice with ABBA. Right, higher magnification of the regions outlined by the dashed-line square (left) in three different behavioural conditions (HC, CTX, FC). Scale bar 100 µm.

Next, we analysed, for each of the three IEGs, which brain regions showed a significant induction in the three behavioural conditions. Brain-wide partial least square analysis (Krishnan et al., 2011; Wheeler et al., 2013) revealed striking differences in the induction of the three IEGs (Fig. 4C, D). Particularly, while Arc and NPAS4 showed increased density in most cortical areas upon exposure to a novel context, cFos increases were less prevalent in the cortex but more prominent in subcortical regions such as amygdalar and hypothalamic areas (Fig. 4C, D). We also investigated whether the three IEGs showed a further induction in animals exposed to CFC as compared to CTX animals. While cFos showed a further increase in a wide set of regions including amygdalar, hypothalamic and brainstem areas, Arc did not exhibit a further induction in animals receiving foot shocks except for a few amygdalar regions and the ventromedial hypothalamus (VMH, Fig. 4D). Notably, NPAS4 displayed a robust induction in the majority of amygdalar regions as well as in prefrontal cortical and retrosplenial cortical divisions (Fig. 4D).

Lastly, we exploited the high atlas registration accuracy ensured by ABBA to create a fine-grained induction map for each IEG and unveil important differences in small subdivisions that may have been masked at a coarser parcellation level (Fig. S6, 7). Indeed, a finer cortical parcellation revealed IEGs induction in CFC compared to CTX animals in specific layers within many cortical divisions including the retrosplenial (RSP), anterior cingulate (ACA), anterior insula (AI), prelimbic (PL) and infralimbic (IL) cortices that could not be detected without a layer specific analysis (Fig. S7). For example, Arc induction was not observed when analysing the PL and RSP at the whole-structure level, but a layer-specific analysis revealed a significant induction in PL layer 6b and RSP layer 2/3. Similarly, a layer 5 specific cFos induction was observed in the ACAd, while NPAS4 induction in the AI was restricted to layer 6 (Fig. S7). These results underscore the importance of dissecting quantifications between specific substructures (such as cortical layers) and therefore emphasise the need for atlas registration tools that ensure high accuracy. Notably, we did not find a global layer specific enrichment in the induction of cFos, Arc or NPAS4, indicating that they behave differently across cortical divisions (Fig. S6, 7).

These results indicate that the brain-wide induction patterns of these three IEGs are largely distinct, with only a small fraction of regions showing comparable induction in all three (Fig 4D, S6,7). These findings suggest that the different IEGs likely reflect different neuronal processes (Mardinly et al., 2016; Hu et al., 2017; Wu et al., 2017; Halawa et al., 2018; Hrvatin et al., 2018). To explore this hypothesis, we investigated whether within each region, the animal’s behavioural state could modulate the probability of neurons to co-express different IEGs. To do this, we measured the number of cells co-labeled by cFos and Arc and normalised by the total number of cFos^+^ and Arc^+^ cells. We did not perform this analysis for NPAS4 as we had observed that the vast majority of NPAS4^+^ neurons co-express cFos. Interestingly, when comparing home cage animals to those exposed to a novel context we found a significant increase in the co-expression probability across most hippocampal and amygdalar subdivisions as well as in a number of cortical structures. In CFC, we observed a further increase in co-expression probability compared to CTX in a more restricted set of areas, including the hippocampal DG, amygdalar BLA, CeA and MEA, claustrum and a few cortical areas such as the mPFC and AI (Fig. 4 E,F), a group of brain areas well-known for their role in fear conditioning (Tovote et al., 2015; Cho et al., 2017). These results suggest that, during different behavioural stimuli, IEG expression may represent heterogeneous activity states and that co-expression of multiple IEGs within the same cells may be associated with more specific activity-related processes. This finding highlights the potential of simultaneous use of multiple markers for IEG-based activity mapping.

## Discussion

Here, we introduce Aligning Big Brains and Atlases (ABBA), a software enabling the precise and efficient registration of high-resolution histological 2D images onto 3D reference atlases. Alongside this, we present BraiAn, a suite for multi-marker automated segmentations, whole-brain optimised statistical analysis and data visualisation. Thanks to the scalability of these tools for multi-marker analysis across large behavioral groups, we compared the brain-wide expression of three widely used IEGs across different behavioural conditions. This analysis revealed significant differences in their distribution and induction, showing these IEGs provide complementary information rather than being equivalent markers of brain activity. In addition, these results suggest that co-expression of IEGs within the same cells may underlie specific activity states and thus represent an option to identify behaviorally relevant neuronal ensembles.

### ABBA/BraiAn a novel toolkit for brain-wide histological analysis

Histological analysis has been a cornerstone in brain research for decades, significantly enhancing our comprehension of this complex organ. Traditionally, researchers have focused on staining specific regions of interest to conduct targeted analysis, however, adopting a more holistic approach that encompasses the entire brain can substantially augment the wealth of information obtained not only in anatomical tracing studies but also in virtually any histological investigations. The recent advent of advanced tissue clearing and imaging techniques like light-sheet microscopy has allowed scaling histological analysis to whole-brain levels (Voigt et al., 2019). However, light-sheet microscopy requires dedicated technology, scaling up to high sample sizes is challenging and it still does not allow efficient detection of multiple markers. To circumvent these limitations, we developed ABBA/BraiAN, a multiplexed, scalable automated pipeline that allows the concomitant quantification of multiple markers across the entire brain using 2D sections. With this, we propose a solution to address the limitations of existing tools and thus facilitate whole-brain histological analysis of large datasets without compromising accuracy. This pipeline includes two open-source freely available packages: 1-ABBA, (Aligning Big Brains and Atlases) that combines machine learning-based technology and image transformation algorithms to flexibly align 2D sections to a 3D reference atlas and 2-BraiAn, (Brain Analysis) to quantify virtually any staining across the whole brain. We designed this pipeline to maximise ease of use, precision, versatility and adaptability. These crucial characteristics are granted by the following features:

**First,** ABBA provides a graphical user interface for all atlas registration steps, making it accessible to researchers with no coding skills. ABBA’s accessibility to a large neuroscience community is further facilitated by a comprehensive set of tutorials and by the inclusion of a simplified installer that drastically simplifies the installation process. Additionally, the active community on image.sc provides a valuable resource for issue resolution and collaborative problem-solving.

**Second,** ABBA excels in compatibility, supporting diverse image data formats through the bio-formats library and seamless integration with the BrainGlobe API (Claudi et al., 2020) for atlas data granting access to numerous 3D atlases including: the adult and developing mouse brain, zebrafish brain, human brain, spinal cord, and rat brain atlases. Noteworthy is its capability to directly read large 2D files from slide scanner microscopes, eliminating the need for lossy conversions. Moreover, the output generated by ABBA is versatile, allowing integration with various downstream tools, with a particular recommendation for the powerful QuPath software.

**Third**, ABBA provides timely and resource-efficient processing of large datasets. Such efficiency stems from the incorporation of well-established tools, including the BigDataViewer ecosystem (Pietzsch et al., 2015) for image display and elastix/transformix (Klein et al., 2010) for registration, while parallelization and asynchronous job handling further contribute to streamlined workflows, ensuring timely and resource-efficient processing of large datasets.

**Fourth**, it combines automatic section positioning, cutting angle adaptation and atlas transformations and at the same time allows manual fine-tuning of each step allowing for ensuring precision and to work with partially damaged sections. User’s online control of ongoing registration steps is provided by real-time visualisation of transformations during the whole automatic registration process. The registration result can be easily stored and shared between experimenters.

**Lastly**, with BraiAn, an integrated suite for quantitative multi-marker image analysis of whole-brain registered datasets, we provide a full package for signal quantification and statistical analysis of whole-brain data (Fig. S2). BraiAn offers a robust platform for large-scale data analysis and visualisation, facilitating data management, atlas navigation at different parcellation levels, custom data aggregation, whole-brain data normalisation, and comprehensive group statistics.

### Differential brain-wide expression of IEGs cFos, Arc and NPAS4

IEG expression represents a versatile and accessible tool for large-scale activity mapping and has been used for decades to assess experience-dependent neuronal activity. A wide variety of IEGs exists, for which different induction patterns and downstream effectors have been identified (Yap and Greenberg, 2018). Despite their well-known molecular divergences however, they are often used interchangeably as a proxy for neuronal activity across brain areas and behavioural stimuli. Here, leveraging ABBA and BraiAn, we performed a systematic analysis and compared the brain-wide distribution of three widely used IEGs, cFos, Arc, and NPAS4 across three different behavioural conditions.

We found that both their relative distribution across brain areas as well as their induction vary substantially. As for their brain-wide distribution, the cortical density of Arc^+^ neurons is significantly higher than that of cFos^+^ while NPAS4^+^ cells are markedly lower. In contrast, subcortical regions such as the pallidum, thalamus, hypothalamus, midbrain, pons, and medulla show high cFos density but undetectable levels of Arc and NPAS4. Interestingly, among these regions, we observed Arc^+^ neurons within the hypothalamic VMH and thalamic RT indicating that their gene expression profiles following activity may differ from their respective hypothalamic and thalamic neighbours.

As for induction of these three IEGs, exposure to a novel context resulted in a consistent increase of Arc and NPAS4 in most cortical areas, while cFos showed less increase in the cortex but was prominently induced in subcortical areas like the amygdala, hypothalamus, and brainstem. In contrast, fear conditioning led to a further increase in cFos in subcortical areas, including the amygdala, hypothalamus, thalamus, and brainstem, while Arc and NPAS4 did not show a widespread increase except in the amygdala, where all IEGs were induced. In addition, co-localization analysis revealed that cFos and Arc co-expression is markedly low but increases following behaviorally salient experiences, while NPAS4 marks a small fraction of the broader cFos^+^ population.

This differential expression of cFos, Arc and NPAS4 may result from different factors. First, expression of these three IEGs may be restricted to specific cell types (and brain areas) expressing different intracellular signalling cascades to translate neuronal activity into IEG induction, which should be addressed by co-labelling IEGs with specific markers for cell identity in future studies. Although the classical view of activity-dependent gene induction posits that calcium-dependent signalling pathways, activated by neuronal stimulation, drive the expression of a common set of IEGs across various cell types, several studies have reported differential IEG induction in distinct neuronal and non-neuronal cell types across a variety of behavioural stimuli and brain areas (Mardinly et al., 2016; Hu et al., 2017; Wu et al., 2017; Halawa et al., 2018; Hrvatin et al., 2018). In addition, the marked subcortical depletion observed for Arc and NPAS4 as opposed to cFos may be mediated by region-specific repressor elements, as reported for the subcortical-enriched NPAS4 repressor factors REST and miR-224 (Bersten et al., 2014).

The second mechanism at the basis of the differential induction patterns could derive from an IEG specific response to different neuronal activity patterns (Mahringer et al., 2022). Indeed, we observed that salient experiences such as exposure to a novel context or to a fear conditioning session induced increased densities of each IEG as well as increased Arc and cFos co-labeling probability, indicating that, differently than background neuronal activity (HC group), experience-dependent neuronal stimulation is reflected in the induction of a shared set of IEGs. Interestingly, single nuclei transcriptomics studies revealed both cell type specific activity-dependent IEG expression as well as IEG expression variability within cell types following epileptogenic stimulation in the cortex (Hu et al., 2017). Similarly, a recent study demonstrated distinct IEGs patterns following neuronal activity in the VTA, varying not only between different cell types but also within the same cell type depending on the source of excitatory inputs (Simon et al., 2024), corroborating the hypothesis that both mechanisms outlined above could be at play.

### Limitations of the study

When interpreting the results, several limitations of our experimental strategy should be considered. First, we report Arc, cFos and NPAS4 brain-wide differential distribution across three different behavioural conditions, but it is important to consider that other stimuli may reveal different expression patterns. For example while we and others observed no NPAS4 in hypothalamic divisions (Bersten et al., 2014; Halawa et al., 2018), its RNA was found in the suprachiasmatic nucleus (SCN) following light stimulation (Xu et al., 2021). A second important consideration is that while for most regions the peak of IEGs activity has been measured around 90 minutes after behavioural exposure, some less studied regions may display different dynamics of IEGs induction and may therefore have been missed in this study. Lastly, our results are based on IEGs protein detection by IHC and are therefore intrinsically bound to antibody efficacy and may not directly reflect mRNA-based IEGs mapping.

### Considerations for the use of IEGs as activity markers

In light of these findings, several key considerations emerge for the future application of IEGs as reliable markers of neuronal activity. First, the choice of IEG is critically dependent on the region of interest. Arc and NPAS4 are more suitable for identifying neuronal activity in cortical areas, whereas cFos is more ubiquitously expressed across various brain regions. In addition, we found that these three IEGs exhibit distinct responses to different activity states. While Arc and NPAS4 may be more sensitive in highlighting activity differences in response to less salient stimuli, they do not differentiate further with increasing behavioural salience. Conversely, cFos is a more versatile marker, capable of detecting a broader range of brain-wide responses. Therefore, the type of experimental condition is paramount when selecting the appropriate IEG. Moreover, with the advent of engram tagging technologies (Reijmers et al., 2007; Garner et al., 2012; Liu et al., 2012; Guenthner et al., 2013; Denny et al., 2014; DeNardo et al., 2019), that are based on IEG promoters, IEG expression has become instrumental in marking and isolating specific neuronal ensembles for subsequent characterization and functional manipulations. Our results thus underscore the importance of the chosen IEG promoter in these studies and highlight the potential of intersectional approaches using a combination of IEGs (Hochgerner et al., 2023). In conclusion, in this study we showed that different IEGs are differentially expressed and differentially induced across the brain and that their combined use provides a more exhaustive description of brain-wide activity states.

## Methods

### Animals

For all experimental procedures male C57BL/6J mice obtained from Charles River were used. Animals were delivered at 6-7 weeks of age and were used for behavioural testing after two weeks of acclimatation. All animals were housed at 22-25° C on a 12 h light-dark cycle (light on 7AM) with water and food ad libitum. Mice were housed in groups of 5 animals and were single housed 2 days before sacrifice. All animals were handled according to protocols and ethical guidelines approved in the Italian animal licence 911/2021-PR approved by the Italian ministry of health.

### Behavioural procedures

Mice were handled 5 times (once per day for 5 days) and habituated to the behavioural room one day prior to behavioural testing. Behavioural experiments were conducted in an isolated room in the morning (between 8:00 to 12:00) and animals were randomly assigned to the different experimental groups. The behavioural apparatus was cleaned between animals with a 70% ethanol solution. Contextual fear conditioning consisted of a 3 min habituation to the conditioning chamber (StartFear Combined system, PanLab, USA) followed by three 2s foot shocks (0.8 mA) with an interval of 28 s. After the shocks, animals were kept in the conditioning chamber for an additional 15 s. Animals belonging to the novel context group were subjected to an identical procedure but did not receive electrical foot-shocks. Animals belonging to the “home cage” group were left undisturbed in the home cage.

### Immunofluorescence

Mice were sacrificed 90 min after the behavioural task. Mice were deeply anaesthetised with ketamine and xylazine (100 mg/kg + 10 mg/kg, intraperitoneally). The brain was removed after transcardiac perfusion (4.0% paraformaldehyde, 1X PBS, pH 7.4) and then postfixed overnight using ice-cold 4% paraformaldehyde solution, after which they were put in sucrose solution for 3 days (30% sucrose, 1X PBS, 4° C). Brains were subsequently frozen at −80°C and 40 um coronal sections were cut with a sliding cryostat (Histo-line, MC 4000). Whole-brain sections were stored at −20 °C in antifreeze solution (sucrose 30%, ethylene glycol 15%, Na-azide 0.02%, PBS). Immunohistochemistry was performed on free floating sections (1 in 5 whole brain series). First, sections were washed 3 times in PBS at RT (10 min each), and subsequently incubated in blocking solution at room temperature (1% PBS, Triton 0.3% and bovine serum albumin, 1%) for 90 minutes under constant shaking. Sections were then incubated in primary antibody solution for two days at 4°C under constant shaking. Two separate immunohistochemistry procedures were performed on parallel series for cFos-Arc and cFos-NPAS4 co-expression experiments. The first contained a mixture of a rabbit anti-Arc antibody (1:1000, Synaptic System, #156 003) and guinea pig anti-cFos antibody (1:1000, Synaptic System #226 008). The second contained a mixture of a rabbit anti NPAS4 antibody (1:1000, Activity Signaling, #AS-AB18A-100) and guinea pig anti-cFos antibody (1:1000, Synaptic System #226 008). Both were dissolved in PBS 1% and Tryton 0.1%. After a 20 min incubation at room temperature, sections were extensively washed in PBST and incubated with Alexa conjugated secondary antibodies: Donkey anti-rabbit 647 (1:1000, Invitrogen, A31573) and Goat anti-guinea pig 568 (1:500, Invitrogen, A11075) diluted in PBST 0.1 % at RT for 2 hours at room temperature under constant shaking. Sections were then washed 3 times with PBS 1% and then mounted on superfrost glass slides (ThermoScientific) with DAPI Fluoromount (Invitrogen #00-4959-52). Images were acquired with a Zeiss Axioscan microscope Z1 equipped with a Hamamatsu Orca Flash 4 camera (2048×2048 pixels, 6.5 µm pixel size) with a 20x/0.8 Plan Apochromat objective. Resulting image’s pixel size is 0.325 µm.

### ABBA design

ABBA has been designed with a focus on user experience and extensibility. It is written in Java and extensively uses the Scijava framework of the ImageJ/Fiji ecosystem. It is built around a model class and a combined view controller class, built with a BigDataViewer graphical user interface (GUI) for responsive display of big image data. It is possible to extend ABBA by implementing other views (a debugging view already exists, and the table included in the BigDataViewer main GUI is a view in itself). It is envisioned to implement additional views: one for Napari and one for Imaris. Importantly, views are optional and ABBA can be used headless either in Java or in Python or with any other language that can execute Java code. During the registration process, nearly all actions (moving slices, registering them) are performed asynchronously by being placed into a queue, each section having its own task queue. This allows to improve the user experience, since the GUI remains reactive even with a heavy CPU load.

The model (i.e. state of the registration) is persisted using a set of human readable json files, packaged into a single zip file. ABBA has remained compatible with all its versions since the start of its development in 2019, a single major change having occurred to reduce and improve error handling during state file opening.

### Image analysis

Whole-brain datasets were first registered to the Allen Brain Atlas (adult mouse brain CCFv3) using ABBA. Slice positioning and angle correction were manually performed. For all sections, Atlas alignments were first performed automatically concatenating affine and spline registrations and subsequently manually refined using BigWarp to maximize precision. For this white matter landmarks and DAPI densities were used. Atlas registration accuracy was checked by at least two independent experimenters. All atlas annotations were imported into QuPath v0.5.1. Quantification of cFos, Arc and NPAS4 was performed on 16-bit grey scale images using BraiAnDetect extension for QuPath software. For each quantified channel, the threshold for positive cell detection (QuPath watershed algorithm) was set at the first peak of the intensity histogram derived from the corresponding image. Histograms were previously smoothed using moving average (window size 15). This was performed to ensure consistent cell detection outcomes on unevenly colour-distributed images. Other automatic segmentation parameters were fine tuned for each IEG in order to minimise false negatives, at the expense of a higher number of false positive detections that were excluded in a second step. This was achieved via QuPath built-in random tree classifier for each IEG. Specifically, the classifiers were trained by an expert user manually labelling all detections in randomly defined areas equally represented across HC, CTX and FC groups. For Arc, due to its visible expression levels difference between cortical and subcortical areas, we trained and applied two different classifiers (one for Isocortex, CTXsp, OLF, CA1, and one for the remaining brain regions). For the RT, classifier corrections were not applied as no false positives detections were present. For all three markers positive cell detection accuracy was verified by at least two trained experts.

For co-localization analysis, we assessed whether cFos^+^ detections contained an Arc^+^ or NPAS4^+^ centroid. To ensure computational performance we used ad-hoc implementation of a bounding volume hierarchy. The whole analysis runs on a representative brain (51 sections, accounting for a total of 102GB) in 36’6’’±7’30’’ (std) on a regular desktop computer (CPU: Intel i9-13900,32 cores @ 5.300GHz, RAM: 63988MiB) accessing the image-sets over a local network.

In order to assure reproducibility and parameters consistency across groups, BraiAnDetect splits script code from a single, user-defined, parameter configuration file which is common between all QuPath projects. Importantly, damaged or mis-aligned tissue portions were excluded from further analysis.

### Whole-brain data analysis

Statistical analysis and whole-brain data visualisation were performed with BraiAnalyse, a custom built Python library enabling fast and accessible analysis of whole-brain data, represented as brain sections, whole-brains or groups. First, we imported the quantification results from QuPath/BraiAnDetect and checked for positive cell density variability within each animal. Thus, for each region, we computed the coefficient of variation 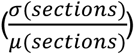 across brain sections, and manually checked those with elevated coefficients. This served as a quality control step for major slip-through, signalling potential problems when the variation of number of detections changed drastically between sections from the same animal. Following this step, we calculated the number of positive cells for each brain region by summing detected cells across all sections. The same was done with the associated structures’ area in mm². All subsequent analysis was based on these raw data reduction, including density (#*IEG*/*mm*²) and similarity index 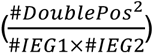 among all (Fig. S2A-C). Based on the chosen metric, we computed statistics for each group using either Pearson correlation, when comparing two markers, or task mean-centered partial least squares analysis (Krishnan et al., 2011). BraiAn’s implementation compares *n* observations structured into *m* groups, across *j* brain regions for which all *n* observations have non-infinite data. Through normalisation, single value decomposition and generalisation of the results by bootstrap & permutation, the PLS provides: 1) a vector <u>*u*</u> of contrasts (or groups’ salience scores) for each comparison suggesting how differentiated a group is from the other *m* − 1; 2) a p-value for each comparison suggesting whether the sampled signal is differentiated from noise; 3) a vector <u>*v*</u> of regions’ salience scores for each comparison suggesting the contribution of each brain region to the groups difference; 4) a projection of the *n* observations onto principal components (for an example see Fig. S2D). We used PLS analysis to compare whole-brain IEGs densities between different behavioural groups, and the resulting contrasts and salience scores were generalised and normalised through permutation and bootstrap testing sampled 10,000 times each. The brain regions’ salience scores are akin to a Z-score, therefore when they are larger than ∼1.96 (i.e., two-tailed p<0.05) the corresponding saliences are considered significantly stable. Finally, when applying PLS between only two groups and when the only latent variable is generalisable, the activity in a brain region is significantly contributing— positively or negatively—to the difference between two groups when its |*salience score*| ≥ ∼1.96.

The study was carried out at multiple levels of granularity in Allen’s ontology: *major divisions*, *summary structures* and *leaf regions* (Wang et al., 2020).

Importantly, regions in damaged tissue portion were excluded from the analysis (see BraiAnDetect) are processed in such a way that all the data of the corresponding substructures and parent regions were discarded as well. Data in regions at the same level of Allen Brain Atlas ontology remain untouched. This is done to avoid considering in the analysis parent structures that had data removed from their subregions. We decided not to apply this strategy to Isocortex’s layer 1, however, since it is often hard to properly align and the number of positive cells in all three markers is negligible.

All abbreviations for brain regions used in this manuscript follow the Allen brain atlas nomenclature.

## Contributions

This study was planned and conceptualised by B.A.S., N.C and C.C. N.C. developed ABBA software C.C. contributed to its QuPath extension and developed BraiAn for the data analysis. L.M.R. and C.C.D. planned and performed IHC experiment as well as behavioural experiment. B.D.B C.C.D. and L.M.R. used ABBA to align experimental slices to the Allen Brain Atlas and results were subsequently analysed by C.C. All authors discussed the results. B.A.S and N.C. wrote the paper, with input from all authors.

## Acknowledgments

We would like to thank Alessandro Bellone, Caterina Giovannini, Alice Jelmoni, Marco Bressi, Stefania Sempreboni, Alessia Marchesin, Edoardo Cavaglià, Cornelius van den Heuvel, Maddalena Pieroni and Lika Ambrosishvili, Lucie Dixsaut, Olivier Burri, Romain Guiet, Olexiy Kochubey for their contribution in histological procedures and ABBA/BraiAn testing. We thank Serena Bovetti, Ludovico Silvestri and Flavio Donato for valuable discussion. We thank Peter Bankhead for his help in developing QuPath and ABBA/BraiAn extensions. We thank Curtis Rueden, Christian Tischer, Adam Tyson, Harry Carey, Guillaume Le Goc, Rémy Dornier, Claire Stoffel, for all their contributions in the many open source parts used by this project (Fiji, brainglobe, QuPath, DeepSlice, etc.), as well as all the existing users of ABBA who have significantly contributed to improve this project by using ABBA and reporting issues. We thank all current and past members of the Silva lab and the FENS–Kavli Network of Excellence for their feedback on the project. **Funding**: the laboratory of BAS is supported by the European Research Council (ERC-2021-STG 101042309), the Fondazione Cariplo (2020-3632), Airalzh (AGYR2021), the Alzheimer’s Association (AARG-22-974392) and the Université Côte d’Azur. Work in the laboratory of J.G. is supported by an ERC Consolidator Grant (CoG 101043457), the Swiss National Science Foundation (310030_219342 and 310030_197752), the Chan Zuckerberg Initiative, and the Synapsis Foundation Switzerland.

## Resource availability

### Lead contact

Further information and requests should be directed to and will be fulfilled by the lead contact, Bianca A. Silva

### Materials availability

This study did not generate new unique materials.

### Data and code availability

- Data reported in this paper are available from the lead contact upon reasonable request.
- All codes generated for this paper are publicly available at: ABBA: go.epfl.ch/abba BraiAn: https://silvalab.codeberg.page/BraiAn
- Any additional information required to reanalyze the data reported in this paper is available from the lead contact upon request.

## Supplementary figures

**Supplementary figure 1:**
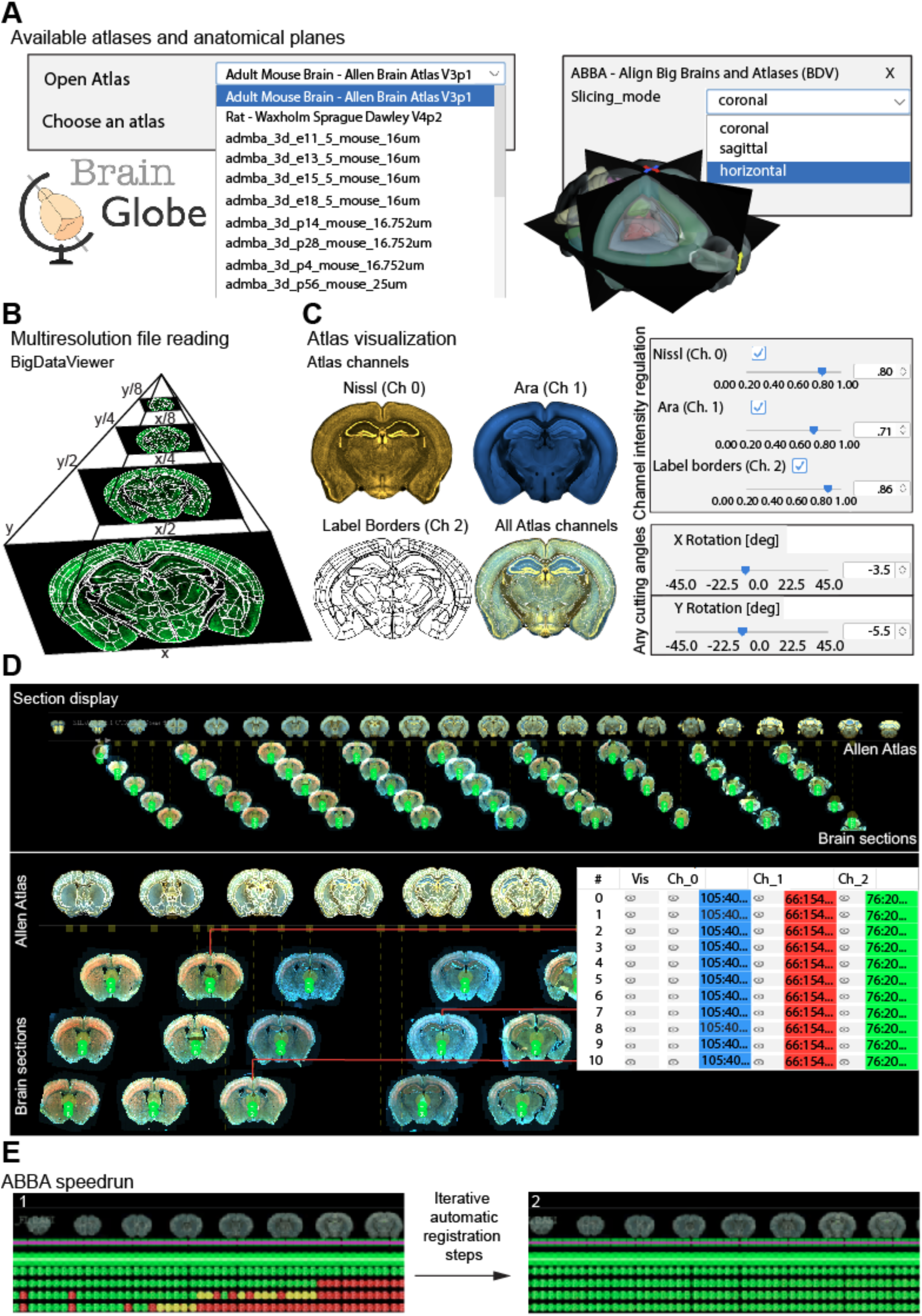
ABBA workflow and capabilities. **A.** Left, the ABBA software interface displaying 10 of the available atlases. Right, ABBA window showing brain representations in coronal, sagittal, and horizontal slicing planes. **B.** BigDataViewer enabling multiresolution data loading using pyramidal data of whole slide images. **C.** Left, ABBA channel visualisation of an atlas slice with corresponding channel display options windows: upper part showing Nissl staining (channel 0), Ara (channel 1), and lower part showing label borders (channel 2) and all atlas channels combined. Right, angle modification window, for virtually depicting any cutting angles, with an example of adjusted cutting angles. **D.** Representative portion of the ABBA GUI for comprehensive visualisation of the experimental dataset. Upper part, Allen Atlas slices (top) and experimental sections (bottom). Lower part, GUI zoom-in including the interactive window for experimental section navigation and custom image display (adjustable brightness and contrast for each image channel) **E.** ABBA registration speedrun. Representative images of GUI during affine and spline registration of selected brain sections to Allen Atlas: green dots denote completed registrations, yellow dots indicate images with registrations in progress and red dots represent images yet to be registered.

**Supplementary figure 2:**
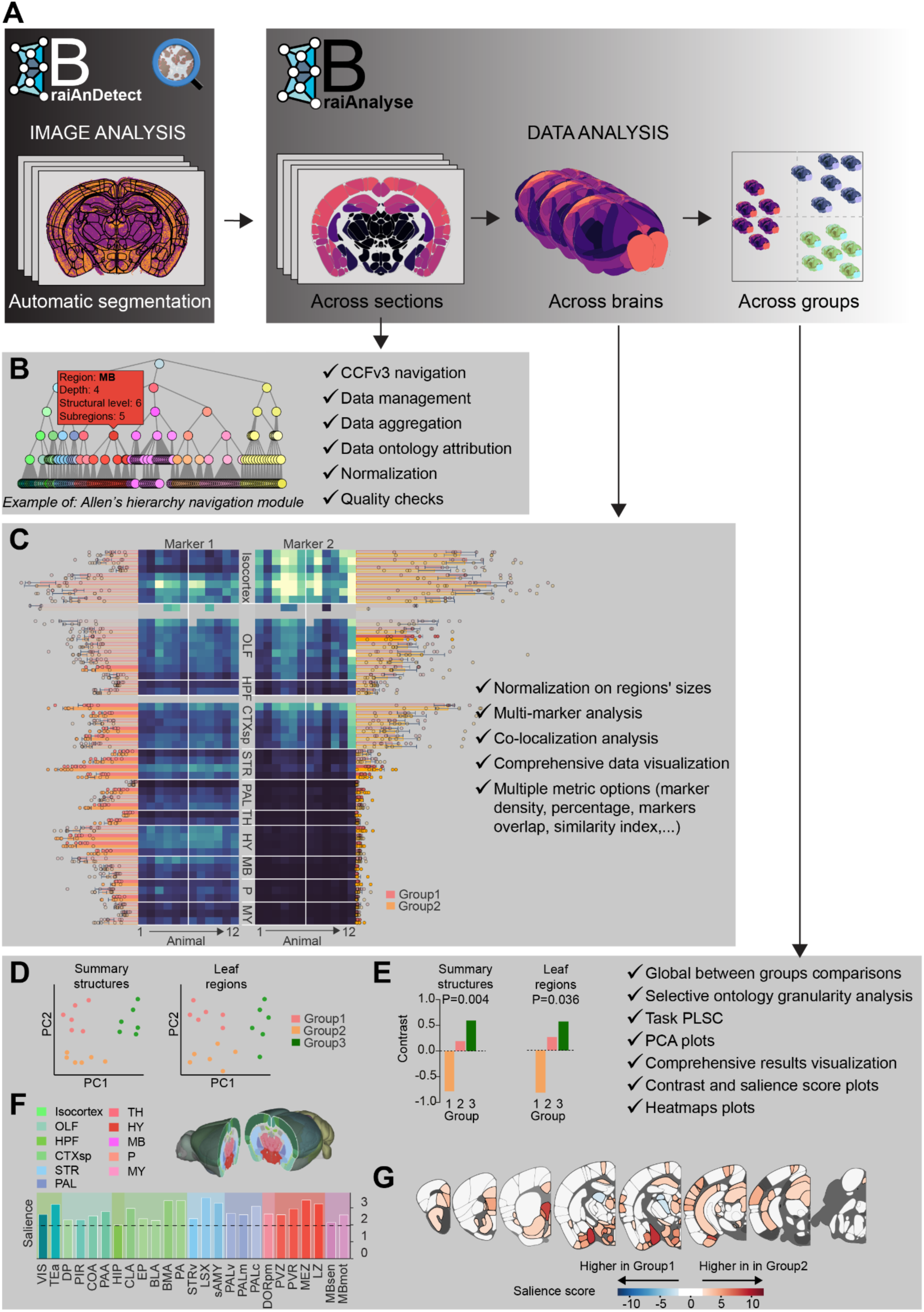
BraiAn workflow and capabilities. **A.** Diagram of BraiAnDetect and BraiAnalyse combined workflow, offering a programmable interface for building a chain of batched image computation and analysis **B.** BriaAn features multiple intuitive data navigation and visualisation options and interacts with Allen Brain Atlas ontology at all stages of data analysis. BraiAn reduces single section data into whole-brain data at the chosen parcellation level featuring various metrics including between sections: sum, mean, standard deviation and coefficient of variation; the latter, along with per-image global marker densities, facilitates the identification of major slip-throughs **C.** All normalised metrics can be visualised in a comprehensive *XMasTree* plot. Here the trunk heatmap displays, for each column, the values of each subject (gray squares indicate regions that were missing in the original dataset), while the branches and the dots display the bar plots for each group in each brain region. Brain regions with solid colours indicate significant salience score from PLS. **D-F.** BraiAnalyse performs between groups whole-brain data statistical comparisons using an optimised PLS algorithm. This implementation yields multiple information about the computed group analysis, including: a PCA style plot (**D**), the groups’ contrasts (**E**) and the normalised salience scores, sorted by major divisions (**F**). **G.** For any analysed metric, BraiAn projects whole-brain data into anatomically represented heatmaps that can be virtually sectioned in any of the 3 axes and angles.

**Supplementary figure 3.**
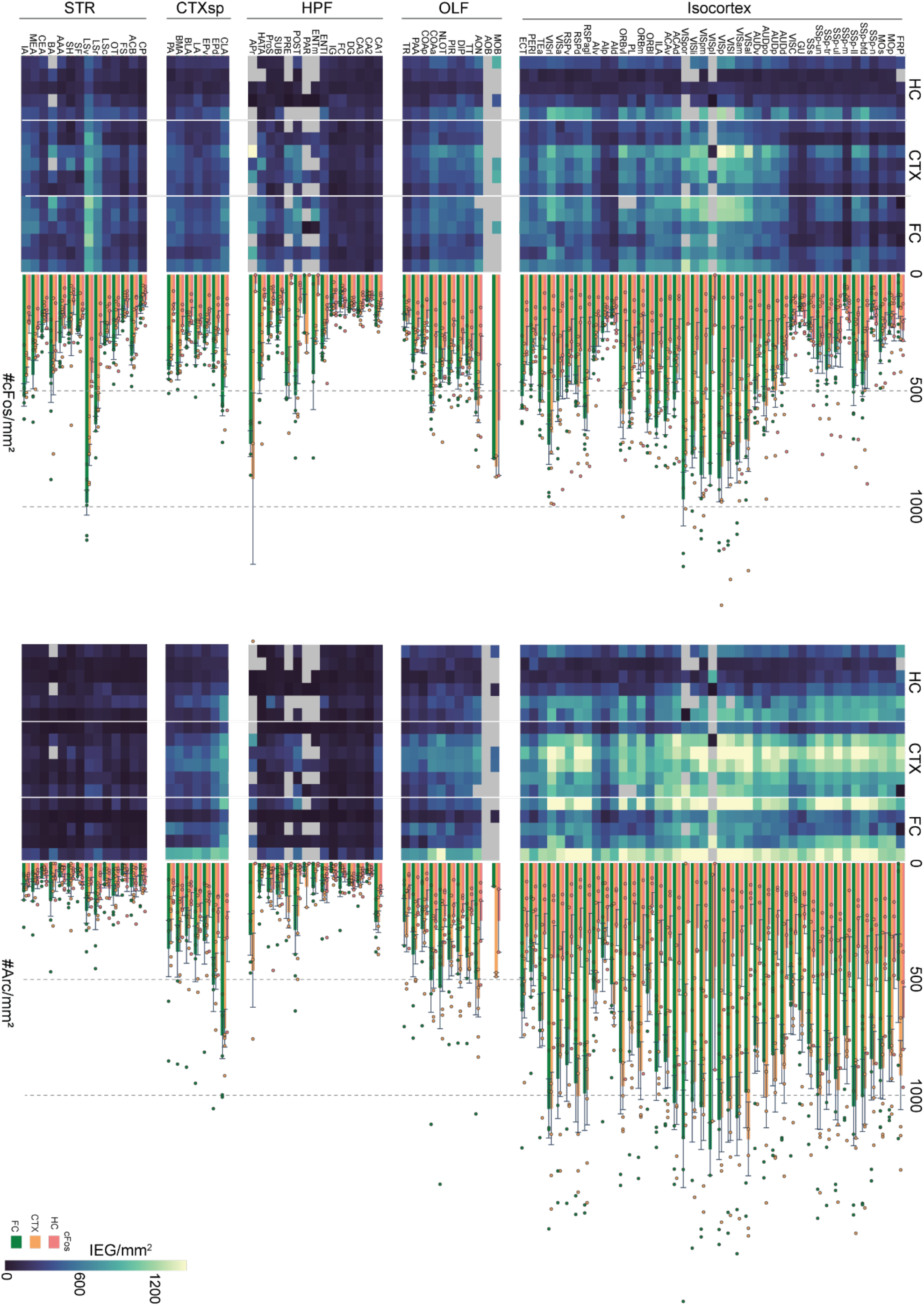
XmasTree plot of brain regions categorised by CCFv3 major divisions: isocortex, olfactory areas (OLF), hippocampal formation (HPF), cortical subplate (CTXsp), striatum (STR), for cFos and Arc for each animal in each behavioural condition (HC, CTX, FC). Top, heatmaps representing densities for each animal (raws) for each region (columns). Gray squares indicate regions that were missing in the images dataset. Bottom, bar plots (Mean ± SEM) of densities in HC, CTX, FC groups across all analysed brain regions.

**Supplementary figure 4.**
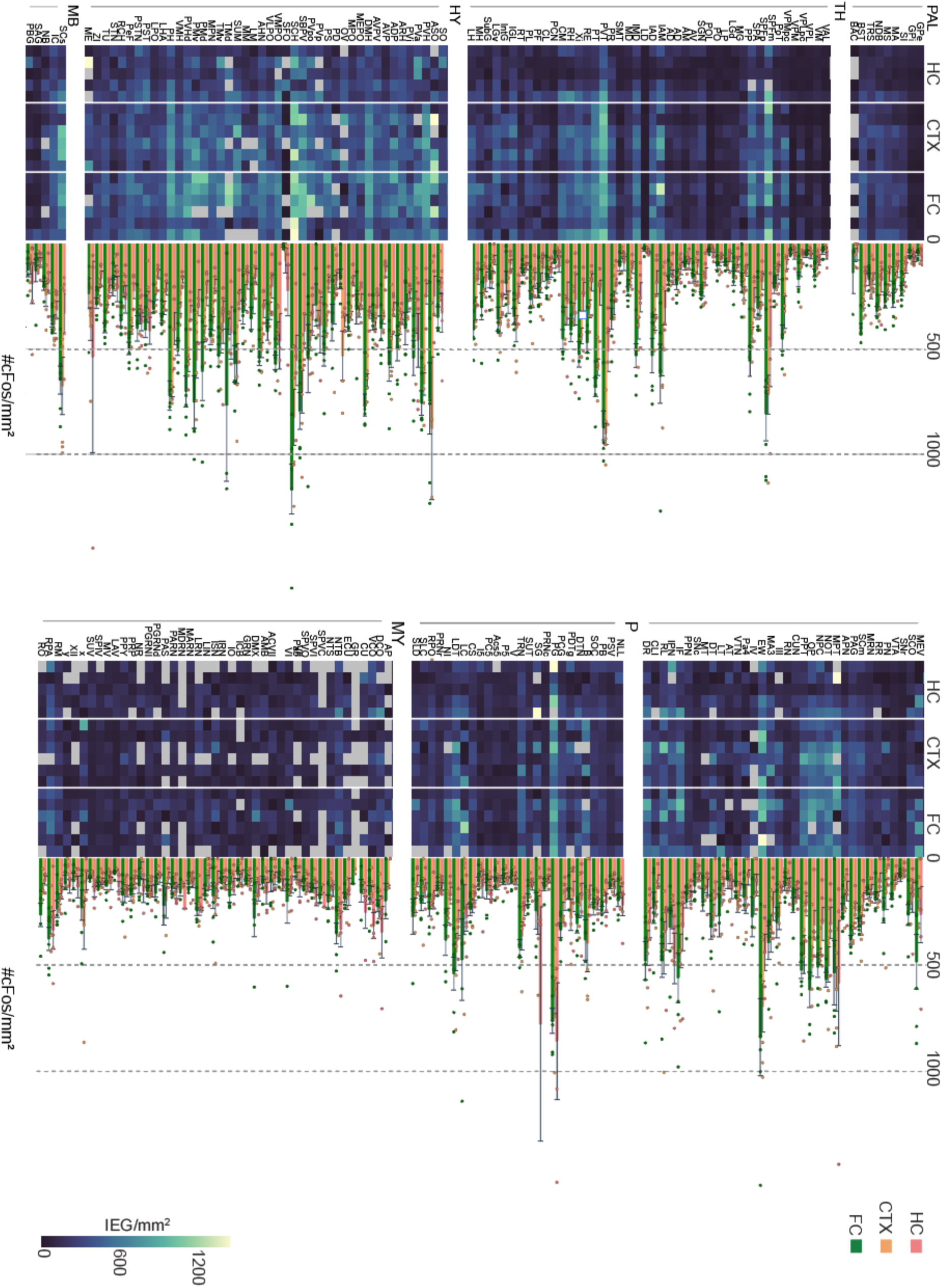
XmasTree plot of brain regions categorised by the following major division: pallidum (PAL), thalamus (TH), hypothalamus (HY), midbrain (MB), pons (P) and medulla (MY) for cFos for each animal in each behavioural conditions (HC, CTX, FC). Top, heatmaps representing densities for each animal (raws) for each region (columns). Gray squares indicate regions that were missing in the original dataset. Bottom, bar plots (Mean ± SEM) of densities in HC, CTX, FC groups across all analysed brain regions.

**Supplementary figure 5:**
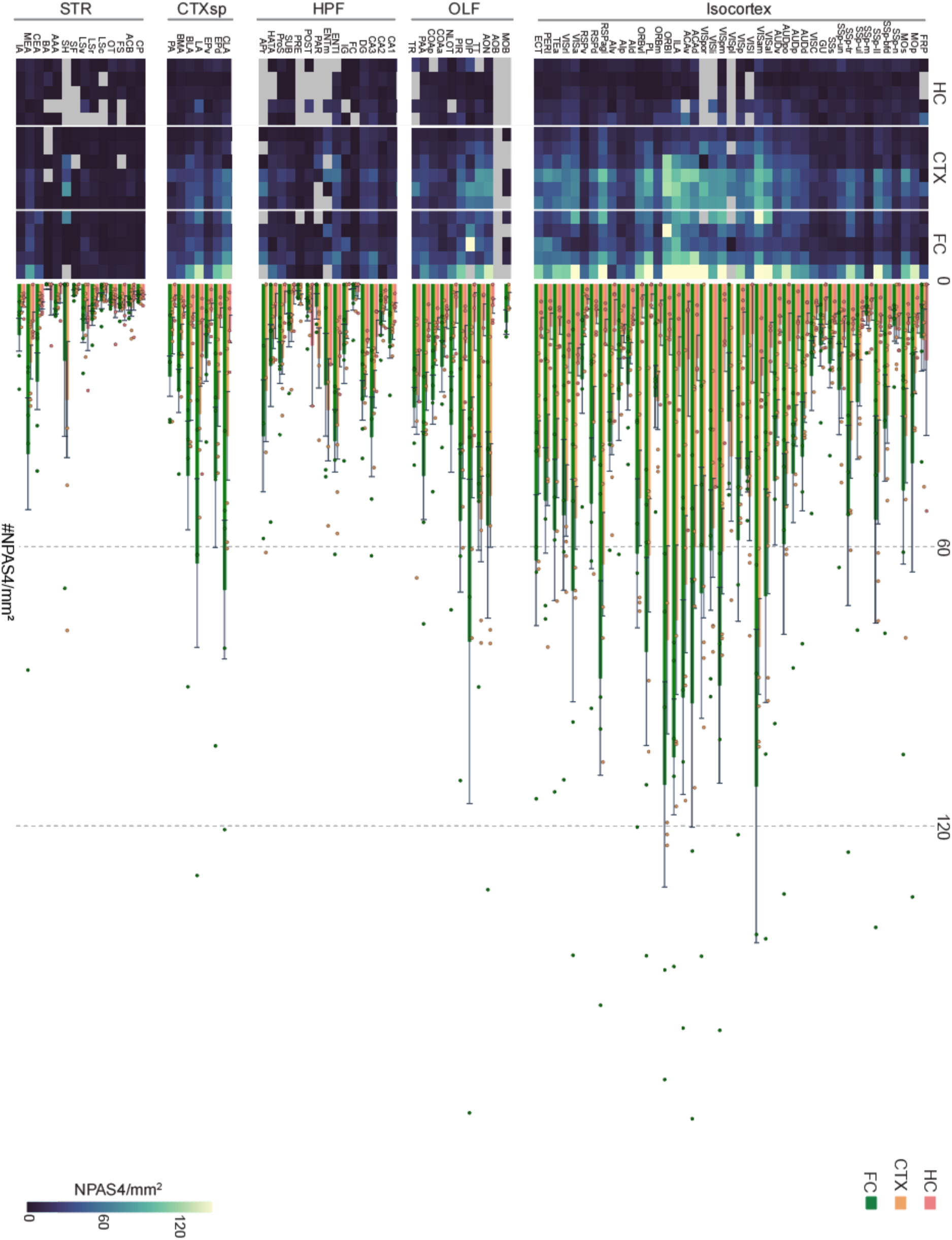
XmasTree plot of all analysed brain regions categorised by the following major divisions: isocortex, olfactory areas (OLF), hippocampal formation (HPF), cortical subplate (CTXsp), striatum (STR), for NPAS4 for each animal in each behavioural conditions (HC, CTX, FC). Top, heatmaps representing densities for each animal (columns) for each region (raws). Gray squares indicate regions that were missing in the original dataset. Bottom, bar plots (Mean ± SEM) of densities in HC, CTX, FC groups across all analysed brain regions.

**Supplementary figure 6:**
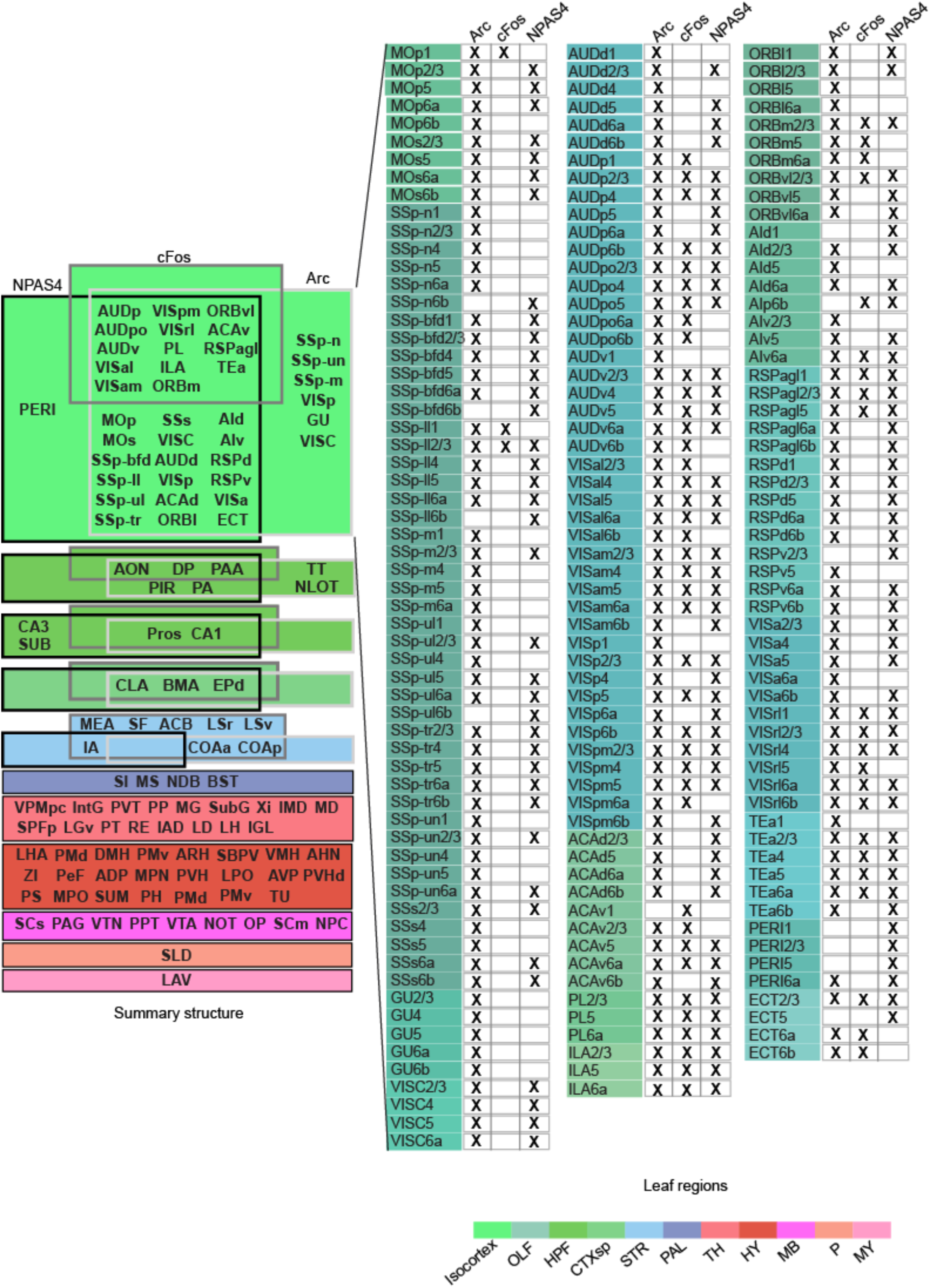
Overview of activity mapping obtained by PLS analysis at different atlas parcellation levels. HC vs CTX for cFos, Arc and NPAS4. Left, list of acronyms of differentially activated regions at the parcellation level of *summary structures* (Wang et al., 2020) as analysed in Fig. 4D; right, acronyms list of differentially activated regions (PLS salience score>1.96) at the parcellation level of *leaf regions* in the isocortex.

**Supplementary figure 7:**
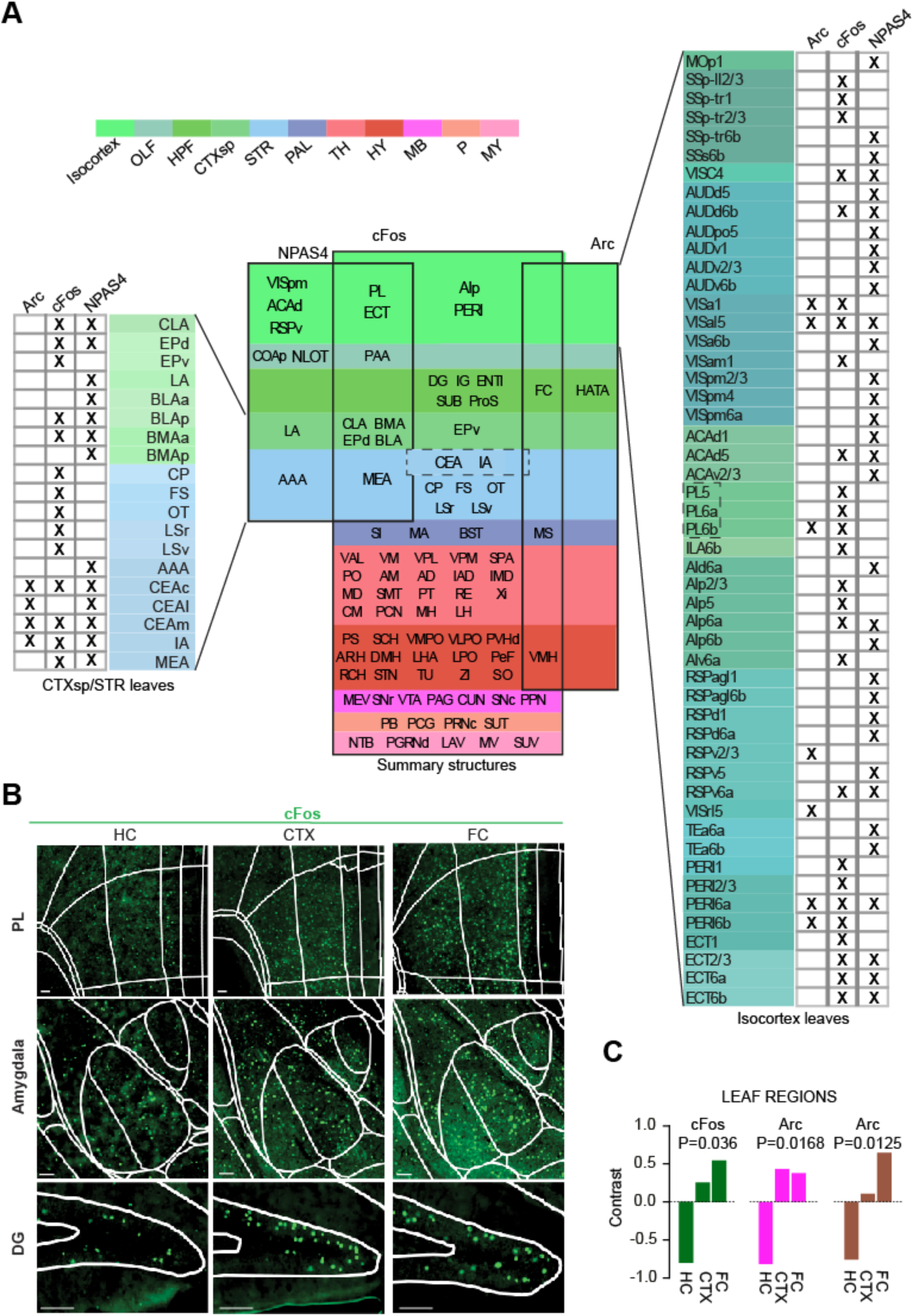
Overview of activity mapping results obtained by PLS analysis at different atlas parcellation levels. CTX vs FC for cFos, Arc and NPAS4. **A.** Center, list of acronyms of differentially activated regions at the parcellation level of “summary structures” (Wang et al., 2020) as analysed in Fig. 4D; left, list of acronyms of differentially activated regions (PLS salience score > 1.96) at the parcellation level of “leaf regions” in the cortical subplate (CTXsp) and in the striatum (STR); right, list of acronyms of differentially activated regions (PLS salience score > 1.96) at the parcellation level of “leaf regions” in the isocortex. **B.** Representative images of cFos immunohistochemistry for HC, CTX and FC in three different sample regions (PL, amygdala and DG). The white reticulate represents the boundaries of Allen CCFv3’s brain regions, after being mapped onto each brain slice with ABBA. Scale bar, 100 µm. **C.** Contrasts of the first latent variable of the mean-centered task PLS analysis between all three groups’ cFos, Arc and NPAS4 density. The latent variables of PLS are generalised with a permutation test (n=10000) and normalised by standard deviation calculated with bootstrap (n=10000). The final score is related to the first left singular vector that maximally differentiates between conditions. Animals: N=6 per group (except for NPAS4 HC, N=5). *Leaf Regions*: cFos: n=395; Arc: n=253, NPAS4: n=213.

